# Unstable Slow Oscillations Couple with Epileptogenic Fast-Rhythm Bistability in Sleep-Related Epilepsy: An SEEG Study

**DOI:** 10.1101/2025.03.25.642592

**Authors:** Gaia Burlando, Chiara Belforte, Felix Siebenhühner, Luca Di Tullio, Lorenzo Chiarella, Vladislav Myrov, Frédéric Zubler, Monica Roascio, Francesco Cardinale, Satu Palva, J Matias Palva, Laura Tassi, Lino Nobili, Gabriele Arnulfo, Sheng H Wang

## Abstract

**Objective:** While slow waves in δ (0.5–4 Hz) characterize NREM sleep, in patients with sleep-related epilepsy, seizures most frequently emerge during NREM stage 2, known to be promoted by δ-band instability. Meanwhile, the epileptogenic zone (EZ) shows localized bistability in β–γ band (15–200 Hz) neuronal oscillations—indicating a catastrophic shift toward seizure. We aim to clarify the mechanistic link between δ-band synchrony and β–γ band bistability in epilepsy.

**Methods:** We studied a cohort of fourteen patients with Sleep Hypermotor Epilepsy (22.3 ± 10.8 years old; 7 males). 7–9-hour stereo-EEG sleep recordings were segmented into 10-minute of uninterrupted, interictal N2 and N3 epochs, and phase synchrony, phase-amplitude coupling (PAC), and bistability were assessed. Canonical correlation was examined to answer whether PAC links δ-phase to β–γ bistability.

**Results:** Compared to non-EZ, the EZ exhibited larger 15–200 Hz bistability along with stronger 2–8 Hz and 15–100 Hz synchrony throughout N2 and N3. Compared to N3, N2 showed stronger PAC between 2–30 Hz phases in the non-EZ and 5–150 Hz amplitudes in the EZ. Canonical correlations between δ-phase modulated PAC and both bistability and synchrony were identified during N2 (*r* = 0.86 and 0.82) and N3 (*r* = 0.84 and 0.80), with the strongest contributors being 2–4 Hz synchrony and bistability in 2–4 Hz and 15–200 Hz bands. Correlations between interictal spikes and canonical covariates of bistability and PAC (*r*^2^ = 0.62 for N2 and 0.56 for N3) validated their relevance to epileptogenicity.

**Significance:** δ-band synchrony and β–γ band bistability are not isolated epileptogenic mechanisms but likely act synergistically, playing a pivotal role in seizure generation through the coupling of δ phases and β–γ amplitudes across large networks, with significant contributions from non-epileptogenic tissues.

**Key points:**

- Strong β–γ bistability in neuronal oscillations localizes the EZ throughout N2 and N3 sleep.
- Elevated δ-band phase synchrony characterizes the EZ and its functional neighbors throughout N2 and N3 sleep.
- ä-band synchrony modulates local β–γ bistability through PAC, with significant contributions from non-EZ tissues.

## Introduction

NREM sleep promotes interictal epileptiform discharges (IEDs) and facilitates seizure activity, whereas REM sleep suppresses them^1–3^. Experimental and clinical studies have demonstrated that thalamo-cortical and cortical sleep oscillations—which drive the spontaneous emergence of slow waves, spindles, and ripples during NREM sleep—can promote paroxysmal epileptic activity^4–8^. Slow waves exhibit an alternation between “up” and “down” states, with up states linked to the generation of spindles and ripples^9–13^. The transition from the down state to the up state—a period of increased probability of synchronous firing—has been identified as a key driver of interictal epileptic activity during NREM sleep^14^. Moreover, several studies have shown that IEDs and seizures are more likely to occur during unstable sleep— particularly in N2—where arousal fluctuations depicted by bursts of EEG slow waves (known as the cyclic alternating pattern) are prevalent^15–20^. In contrast, IEDs and seizures occur less frequently during stable sleep (N3) when slow-wave activity peaks. These findings challenge the traditional view that slow waves directly promote epileptic activity. Instead, epileptogenicity may be linked to large-scale network dynamics mediated by slow waves with prominent waxing-and-waning dynamics, a hallmark of *bistability* in complex systems, such as the brain, operating near criticality^21,22^.

The *brain criticality* hypothesis proposes that the brain gains functional benefits from operating near a critical transition between asynchrony and hypersynchrony^23^. In this context, epilepsy is associated with excessive synchronization with an underlying shift towards a supercritical regime^24^. *Bistability* describes the brain’s tendency to alternate between periods of low and high network activity (or synchrony), where internal feedback mechanisms make these shifts more likely to repeat and amplify over time. Across a wide range of computational models^21,25–28^, strong bistability is thought to precede catastrophic events, where violent activity, such as hypersynchronous seizure activity, can suddenly emerge from normal or quiescent states (Fig. 1A). Recent studies have shown that the epileptogenic zone (EZ) exhibits pronounced β– γ (15–200 Hz) bistability^21,22^, and that this phenomenon becomes even more pronounced in sleep-related epilepsies during NREM sleep, when high-amplitude β–γ bursts occur more frequently than during wakefulness^29^.

**Fig 1.**
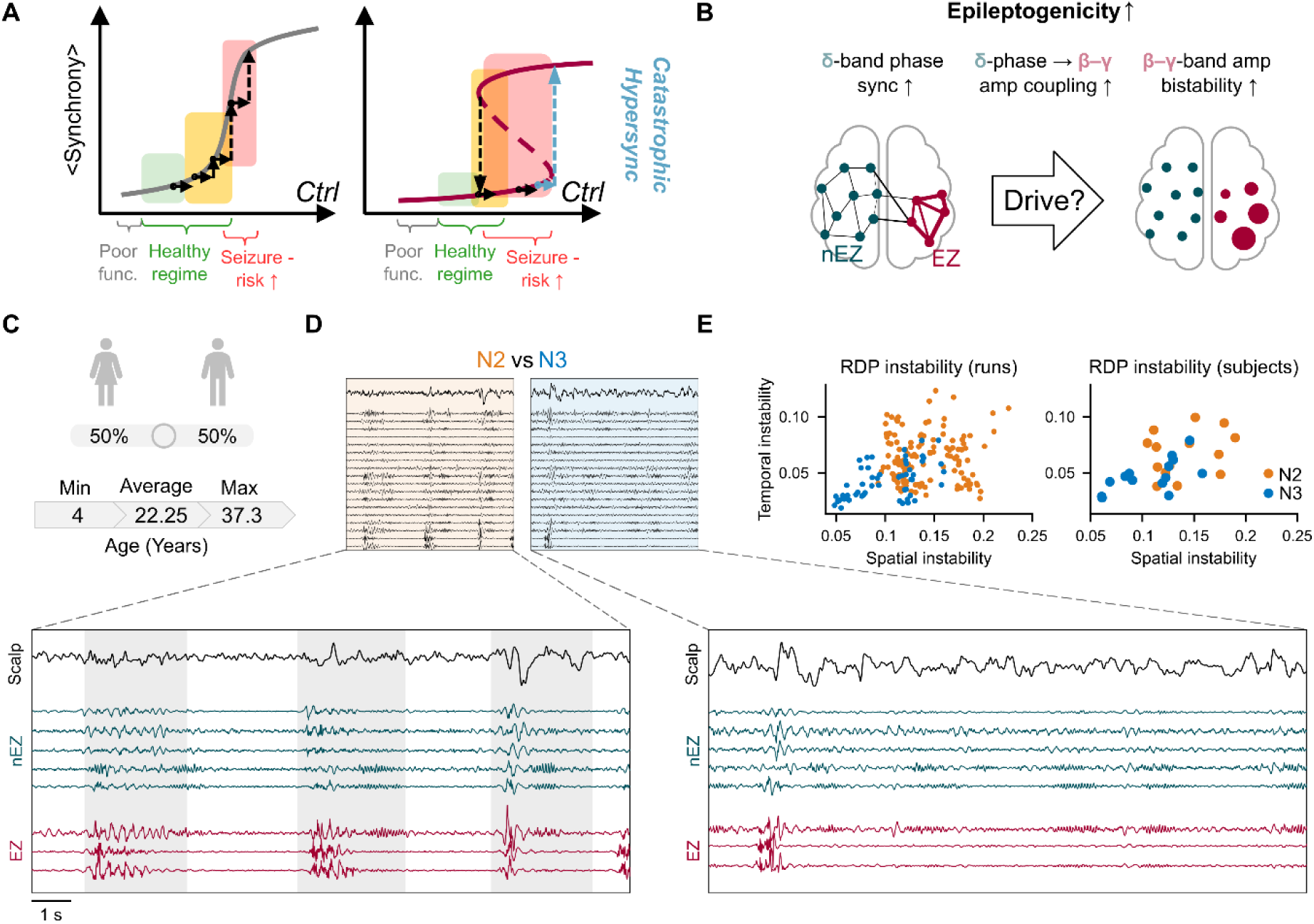
Study schematics. **(A)** As excitability (ctrl) increases, the nEZ exhibits classic criticality with unimodal mean synchrony (left), whereas the EZ displays strong bistability (right) mean synchrony in neuronal oscillations. **(B)** The EZ shows strong δ-band synchrony (left) and β–γ band bistability (right). *Hypothesis*: a mechanistic link between δ phase and β-γ bistability (center). **(C)** Subjects. **(D)** Exemplary bipolar referenced 15-second traces. Only a subset of channels is shown. **(E)** Replication of δ-band instability in N2 and N3 sleep. Epoch-level (left) and individual-level (right) spatial and temporal δ-instability during N2 (orange) and N3 (blue).

**Fig 2.**
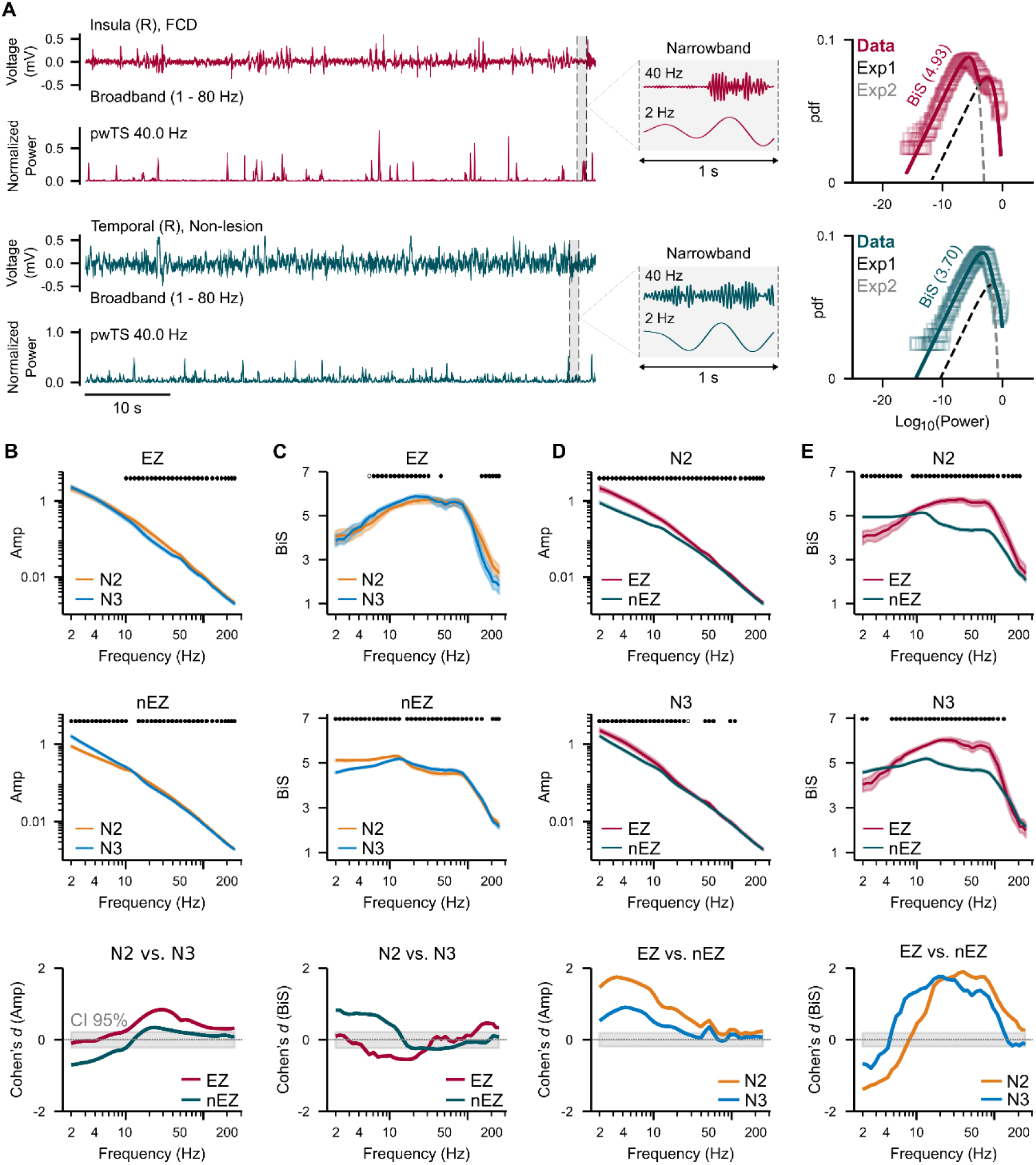
The EZ exhibits stronger β–γ bistability in local oscillations. **(A)** Left: bipolar-referenced broadband (BB) and its 40 Hz narrowband power time series (pwTS) from an EZ (top) and a nearby nEZ (bottom) channel from a N2 epoch. The insets show a 1-second segment of the 40-Hz and 2-Hz narrowband signals. A bi-exponential fit of the probability density function (pdf, top right) yields a larger bistability estimate (BiS) than a unimodal fit. **(B)** Mean amplitude spectrum for EZ (top), nEZ (middle), and effect size of the difference between N2 and N3 (bottom). **(C)** The same as in **(B)** for bistability. **(D)** Mean amplitude spectrum of EZ and nEZ during N2 (top), N3 (middle), and effect size of the difference between EZ and nEZ (bottom). **(E)** The same as in **(D)** for bistability. Sample size in **(B–E)**: (*N*_EZ_=93; *N*_nEZ_=1046 channels); in **(B–E),** markers indicate difference (Wilcoxon rank-sum test, *p* < 0.05, corrected with Benjamini-Hochberg procedure); shaded areas around the mean curves represent the variability computed from 100 bootstraps (95th percentile); gray shaded areas indicate 2.5% and 97.5%-tile of 10,000 surrogates. Effect sizes of Cohen’s *d*: small (0.2), medium (0.5), and large (≥0.8).

In physiological conditions, large-scale neuronal communication is spontaneously orchestrated by a network of slowly oscillating, loosely-coupled neuronal populations^30,31^. In epileptogenic networks, this physiological coordinating mechanism appears maladaptively engaged within the EZ^32^ which is characterized by elevated δ-band (0.5–4 Hz) phase synchrony and cross-frequency phase–amplitude coupling (PAC) between the phase of δ–θ (0.5–8 Hz) oscillations and the amplitudes of γ (40– 80 Hz) or high-frequency oscillations (>80 Hz); importantly, the phase of slow oscillations predicts seizure-onset^33^. Collectively, strong bistability and elevated synchrony likely co-occur as signatures of the same epileptogenic mechanism at local and larger spatial scales. However, prior studies reported focused only weak to moderate correlations (maximum *r²* ≈ 0.25) between narrow-band synchrony and bistability^22,24,34^, thus questioning a strong mechanistic link between them in discrete frequency bands. Notably, these correlations were assessed primarily in interictal, awake resting-state recordings. We therefore hypothesized that δ-synchronized populations regulate local excitability via PAC in sleep, and that epilepsy hijacks this mechanism. Here, during NREM, we test whether β–γ bistability varies in synchrony with unstable δ-phase via a PAC mechanism^22,24,34^ (Fig. 1B).

As a precaution before a wider analyses spanning different etiologies, we examined a curated dataset of sleep SEEG recordings from fourteen patients with Sleep Hyper-motor Epilepsy (SHE) and type 2 Focal Cortical Dysplasia (FCD) (Fig. 1C). We focused on this cohort because SHE with FCD2 represents a prototypical model of sleep-related epilepsy, with a well-characterized substrate that markedly increases seizure risk during sleep^7,35^.

## Materials and Methods

### Overview

After standardized preprocessing, continuous sleep-SEEG recordings were segmented into 10-minute epochs of clean, uninterrupted NREM sleep stages 2 and 3 (N2, N3). Analyses proceeded in three steps. (1) We compared EZ and nEZ activity within and across N2 and N3, focusing on oscillation amplitudes, bistability, and phase synchrony, with the expectation that δ-band bistability characterizes unstable sleep^19^(Fig. 1D–E) and that elevated δ-synchrony and strong β–γ bistability jointly characterize the EZ. (2) Cross-frequency PAC analysis was next conducted to assess EZ-specific and nEZ-specific connectivity differences between N2 and N3. (3) Canonical correlation analysis (CCA) was used to test whether δ-phase couples strongly to β–γ oscillations, showing elevated bistability, and whether δ-driven cross-frequency PAC network structure also correlates with δ-phase synchrony networks. Consistent results across steps 1 and 2 would replicate previous findings and provide validated interim support, while step 3 would yield novel converging evidence for a cross-frequency mechanism linking synchrony and critical bistability in sleep-related epilepsy.

### Subjects and recordings

The patients who underwent SEEG study in the “Claudio Munari” Center for Epilepsy Surgery (Milan, Italy) were retrospectively screened for this study. The selection criteria were: 1) presence of type 2 FCD lesion confirmed by postsurgical histopathological analysis; 2) Presence of recordings from scalp EEG electrodes, electrooculogram, and submental electromyography electrodes; 3) At least one contact located within or at the immediate border of the FCD, identified based on neuroradiological findings and/or a typical intracerebral EEG pattern; 4) Presence of interictal spikes both within and outside the seizure onset zone^36^; and 5) Complete seizure freedom with a minimum of two years of post-surgical follow-up (Engel class 1A^37^), confirming the localization and complete resection of the EZ. We analyzed SEEG data from fourteen subjects who met these criteria (22.2 ± 10.8 years old, 7 male). SEEG was recorded using Microdeep electrodes D08 (Dixi Medical) or Depth Electrodes Range 2069 (Alcis, France), comprising 5–18 contacts spaced 1.5 mm apart, using a Neurofax EEG-1100 system (Nihon Kohden, Tokyo, Japan) with 1 kHz sampling and 16-bit resolution. Recordings were hardware-filtered with a 500 Hz low-pass; no notch or additional filters were applied. The brain regions explored during SEEG implantation for each subject are reported in the Supplementary Tab. 2. This study was conducted in accordance with the recommendations of the Ethics Commission of Ospedale Niguarda (ID: 939). All patients provided written informed consent for the retrospective analysis of their data, in compliance with the Declaration of Helsinki.

### Clinical identification of the EZ

The EZ is the minimum amount of cortex that must be resected, inactivated, or disconnected to achieve long-term seizure freedom^38^. In this study, the EZ was identified by expert epileptologists as the seizure onset zone through concordance with electro-anatomic-clinical observations, later confirmed by surgical outcome and histology after seizure resection. The EZ for each subject, together with the type of cortical dysplasia confirmed by histopathology, is reported in Supplementary Tab. 2.

### SEEG preprocessing

In these fourteen subjects, 377 out of 1872 SEEG contacts (20.14% per subject) were excluded due to significant artifacts or their location in the white matter. After SEEG implantation, MRI co-registered to post-implant CT was used to classify contacts as cortical or white matter, and the channels were marked in the clinical annotation. For this retrospective analysis, the original MRI files were no longer available, preventing atlas-based mapping; therefore, we relied on the original MRI-based annotations, which were consistent with the lower-amplitude physiology characteristic of white matter. Line noise and its harmonics were removed using a filter bank of notch filters with 1 Hz band-stop width. We employed a bipolar referencing scheme by pairing neighboring contacts and excluding a common reference, yielding N=1139 bipolar-referenced channels, which were used for all subsequent analyses. These were then categorized based on clinically identified FCD: channels with both contacts entirely within the dysplasia were designated as EZ channels (N=93), while those with both contacts entirely within healthy tissue were classified as nEZ channels (N=1046). Channels that did not meet these criteria (*i.e.*, bipolar pairs with one contact within and the other outside the dysplasia) were excluded (Tab. 1).

**Tab. 1.**

| Subject | 01 | 02 | 03 | 04 | 05 | 06 | 07 | 08 | 09 | 10 | 11 | 12 | 13 | 14 | Tot. |
| --- | --- | --- | --- | --- | --- | --- | --- | --- | --- | --- | --- | --- | --- | --- | --- |
| N2 epochs | 7 | 8 | 14 | 10 | 21 | 4 | 8 | 7 | 8 | 11 | 8 | 4 | 12 | 12 | 134 |
| N3 epochs | 5 | 7 | 8 | 3 | 2 | 2 | 2 | 3 | 7 | 5 | 4 | 2 | 4 | 2 | 56 |
| EZ channels | 14 | 2 | 9 | 7 | 11 | 7 | 3 | 4 | 6 | 9 | 3 | 4 | 8 | 6 | 93 |
| nEZ channels | 53 | 67 | 66 | 88 | 55 | 73 | 76 | 116 | 115 | 69 | 74 | 98 | 53 | 43 | 1046 |

**Dataset description.** Individual number of recorded epochs for N2 and N3, number of channels recording from EZ and nEZ, and cumulative total elements.

### Obtaining interictal N2 and N3 epochs

The SleepSEEG Matlab toolbox was used to perform semi-supervised automatic sleep staging on 7–9 hours of uninterrupted SEEG recordings^39^. From this staging, all possible 10-minute epochs of N2 and N3 were extracted. Epochs were retained only if they were artifact-free, belonged to the correct sleep stage, and did not contain seizure activity, following visual inspection and validation by experts. A total of 134 and 56 epochs were obtained for N2 and N3, respectively. On average, the temporal distance between the onsets of any two selected blocks within a patient was 46.8 minutes for N2 and 61.0 minutes for N3.

### Epileptic window detection

Interictal spikes are characterized by high-amplitude, fast activity with widespread spatial diffusion in SEEG data, which, if left untreated, would inflate synchrony, bistability, and PAC assessments. We identified and removed time windows containing spikes, following the approach used in our previous study^40^. These windows were defined as 500 ms non-overlapping intervals during which at least 10% of channels exhibited abnormal sharp peaks—amplitude envelope peaks exceeding five times the standard deviation of the channel mean amplitude—in more than half of the 50 frequency bands (from 2 to 450 Hz). This procedure ensured that the entire set of channels was retained, and only contaminated time windows were removed. On average, 12.08 ± 12.5 such windows were excluded for each 10-minute segment.

### Filtering and assessing neuronal features

We applied a narrow-band Morlet wavelet transform (*m* = 7.5) to the SEEG epochs to obtain time–frequency representations^41^, using 40 logarithmically spaced frequencies between 2 and 250 Hz. Feature extraction was performed in the same manner as in our previous studies^22,40,42^, with formal definitions provided in the Supplementary Methods. Briefly, we assessed relative δ-power as well as narrow-band amplitudes and bistability for narrow-band neuronal oscillations. For large-scale connectivity, we evaluated both 1:1 phase synchrony and PAC using the phase-locking value.

### Canonical correlation analysis (CCA)

CCA was used to first examine relationships between δ-modulated outward-PAC and synchrony as a validation, and between δ-modulated inward-PAC and bistability for the hypothesis testing (Fig. 5A; see Supplementary for formal definitions). Outward-PAC describes how phase of slow waves (*θ_LF_*) in one region correlates with fast activity (*A_HF_*) in other regions, whereas inward-PAC reflects how slow waves from the surrounding network correlate fast activity within that region (Fig. 5A). Rather than independently computing correlations between each pair of bistability (*A*) and synchrony (*B*) estimates across multiple narrow-band frequencies^21,22,24^, CCA identifies *weighted combinations* of narrow-band components in each set—canonical variates (*Z_A_* and *Z_B_*; Eq. 10)—that are most strongly correlated (Fig. 5B). This approach provides a powerful way to test whether bistability and inward-PAC, as well as synchrony and outward-PAC, are strongly correlated at the multivariate level. The corresponding loading (*u, v*), *i.e.*, the weights, indicates the contribution of each frequency to the observed correlation. For example, if both bistability and δ-band correlated with inward-PAC showing strong loadings in the β–γ range, this would indicate that regions whose β–γ amplitudes are strongly correlated with δ-phase also exhibit β–γ bistability, thereby supporting a strong mechanistic relationship between large-scale δ-phase and local β–γ bistability.

To test the hypothesis, we focused on the δ-phase portion (Fig. 5C, top left) of the full cross-frequency PAC design matrix (illustrated in Fig. 4). As each subject had a different number of N2 and N3 epochs, individual PAC connectivity matrices (*e.g.*, 1→80 Hz) were first averaged within subjects so that one PAC matrix represents the mean coupling between a δ-frequency (1 Hz, Fig. 5A, left) and the amplitude of a faster frequency (80 Hz corresponding to ‘ratio of 80’ for 1 Hz) for a N2 or N3 epoch. Inward- and outward-PAC connectivity are scalar vectors obtained by averaging each PAC connectivity matrix column-wise and row-wise, respectively, for each LH–HF pair (Fig. 5C). Note that ‘inward’ and ‘outward’ here refer to the network structure of the PAC graphs based on LF→HF connectivity, without implying causal relationships between the slow and fast modes. Pooling the individual inward- and outward-PAC resulted in corresponding two-dimensional feature arrays, which were then subjected to CCA analysis. Eigenvector centrality (EVC) was used as phase synchrony feature as it demonstrates stronger differentiation power for the EZ (Fig. 3C).

**Fig 3.**
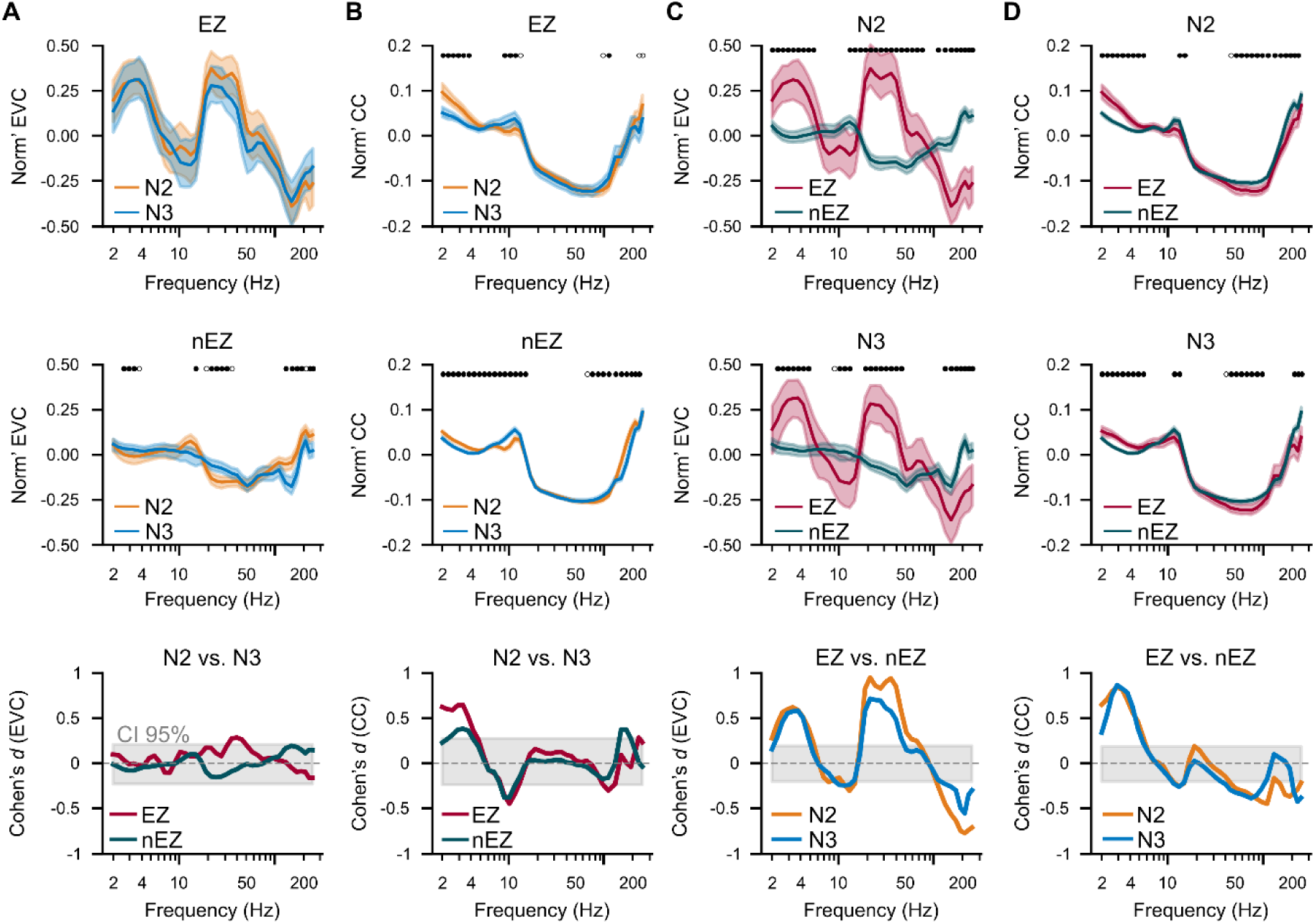
The EZ exhibits stronger large-scale synchrony than the nEZ in both N2 and N3. **(A)** Normalized eigenvector centrality (EVC) during N2 and N3 for the EZ (top), nEZ (middle), and the effect size of the differences between N2 and N3 (bottom). **(B)** The same as in **(A)** for clustering coefficient (CC). **(C)** Normalized EVC of EZ and nEZ during N2 (top) and N3 (middle), and the effect size of the differences between EZ and nEZ during N2 and N3 (bottom). **(D)** The same as in **(C)** for CC. Sample size in **(A– D)**: (*N*_EZ_=93; *N*_nEZ_=1046 channels). Narrow-band features were normalized as x_norm_ = (x_i_ – median(X)) / max(X – median(X)), where X is the feature vector across channels. Markers in **(A–D)** indicate difference (Wilcoxon rank-sum test, *p* < 0.05, FDR corrected with Benjamini-Hochberg procedure); shaded areas around the mean curves represent the variability computed from 100 bootstraps (95th percentile); gray shaded areas in bottom row panels indicate 2.5% and 97.5%-tile of 10’000 surrogates. Cohen classified effect sizes as small (*d* = 0.2), medium (*d* = 0.5), and large (*d* ≥ 0.8).

**Fig 4.**
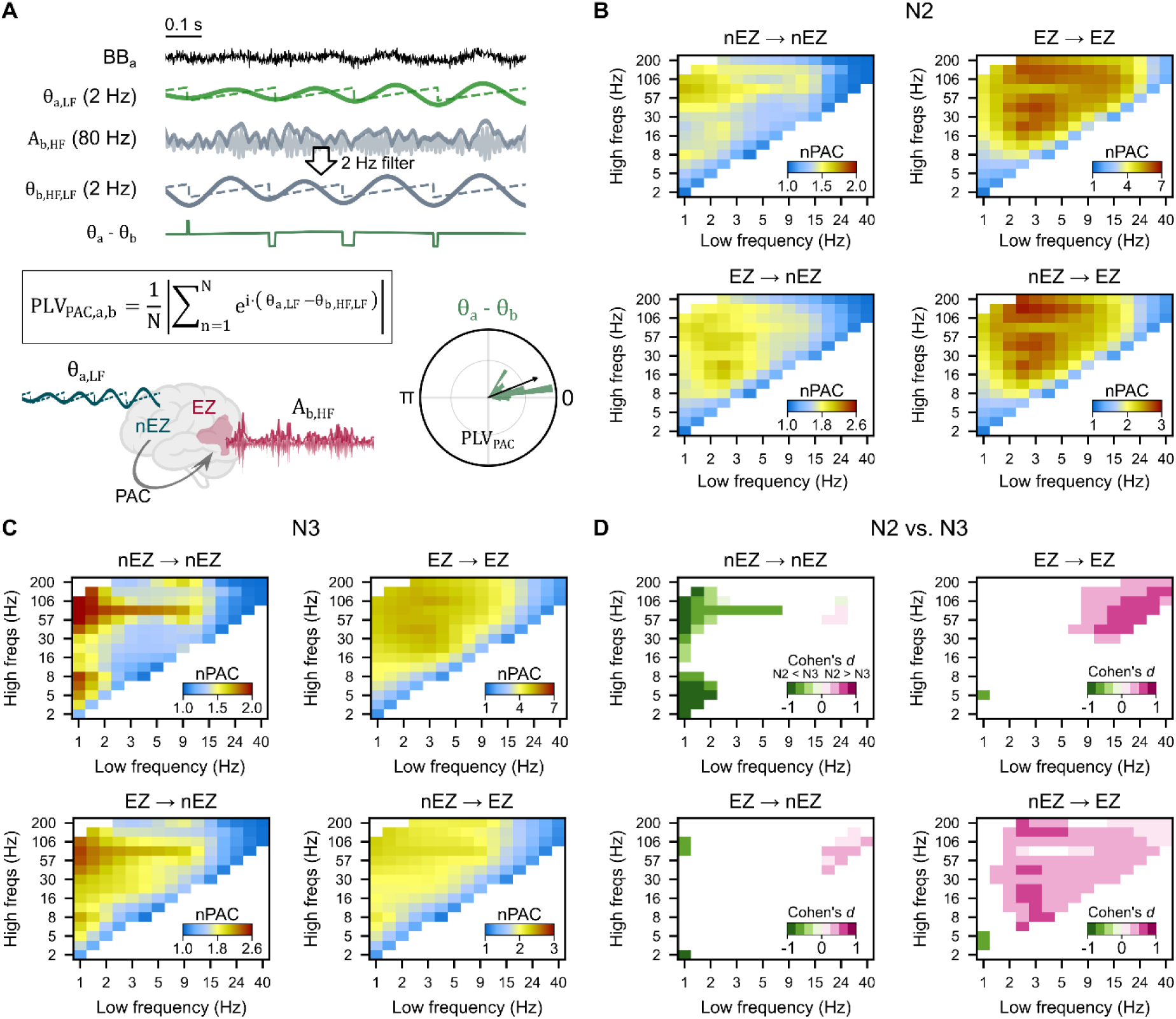
N2 is characterized by stronger coupling between δ (2–5 Hz) phase in the nEZ and amplitudes of fast oscillations in the EZ. **(A)** PAC computation illustrated. **(B–C)** Cohort average of normalized PAC (nPAC) for **(B)** N2 and **(C)** N3; nPAC = PLV_PAC,observed_/PLV_PAC,surrogate_ (N_surrogate_ = 100) so that nPAC > 1 indicates PAC above the null hypothesis level. **(D)** Effect size of the difference between N2 and N3 sleep stages. Only effect sizes for which the differences between N2 and N3 were statistically significant (*p* < 0.05, corrected with Benjamini-Hochberg procedure) are presented. In the cross-frequency PAC design matrix, the x-axis represents low frequencies, and the y-axis represents high frequencies. Each pixel in the colormap represents a specific slow–fast frequency pair of PAC connectivity.

We averaged the canonical variates (CVs) of channel-wise PAC, bistability, and synchrony estimates (samples in Fig. 5D–E scatter plots) within each subject and examined correlations between these coarse-grained neuronal features and the number of spikes observed in the corresponding N2 and N3 epochs. Data from fourteen subjects, comprising a total of 1,139 SEEG bipolar-referenced channels, were used for this analysis. The CCA was carried out using the Python *scikit-learn* module^43^.

**Fig 5.**
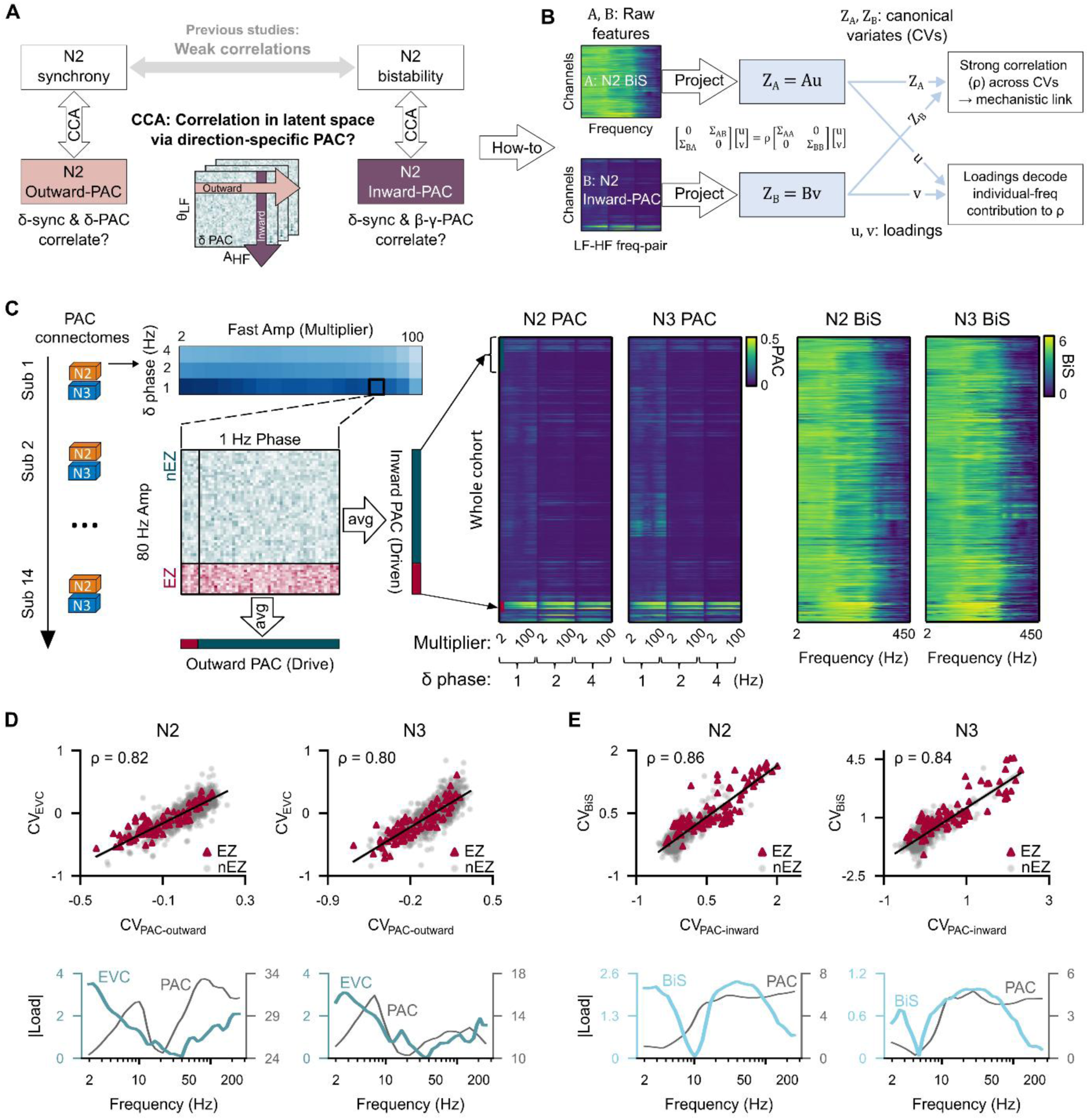
Cross-frequency link between δ-modulated inward-PAC and β–γ bistability. **(A)** Design for canonical correlation analysis (CCA) of N2 illustrated. **(B)** Canonical variates (CVs) and loading illustrated (formal definition in Supplementary). **(C)** Feature data for CCA. **(D)** Top row: Pearson’s correlations between CV of outward-PAC and eigenvector centrality (EVC) (Sample: *N*_EZ_=93; *N*_nEZ_=1046 pooled from 14 subjects). Bottom row: loadings of outward-PAC and EVC showing the contribution of each frequency to the observed correlation. **(E)** The same as **(D)** but for inward-PAC and bistability.

## Results

### Replication of greater δ-band instability in N2 compared with N3 sleep

We compared δ-band instability between N2 and N3 using the Wilcoxon rank-sum test (α = 0.05), considering both spatial instability (across channels) and temporal instability (across time for a channel). When treating each 10-minute run as an independent observation (134 runs for N2, 56 runs for N3), we found that N2 compared to N3 shows significantly higher spatial (*p* < 6.5 × 10^-9^) and temporal instability (*p* < 8.8 × 10^-10^) (left, Fig. 1E). However, when averaging δ-band instability within subjects, only temporal instability remained significantly higher in N2 compared to N3 (*p* < 0.013), while spatial instability did not reach statistical significance (*p* > 0.054) (right, Fig. 1E).

### Strong δ-band amplitudes and β–γ band bistability characterize the EZ

We next examined whether N2 and N3 sleep can be differentiated by amplitudes and bistability across narrow-band oscillations. Within nEZ, 2–10 Hz amplitudes were higher in N3 than N2 (Fig. 2B; *p* < 0.05, Wilcoxon rank sum test, multiple comparison corrected with BH procedure; negative Cohen’s *d* indicates N3 > N2), whereas no similar stage-dependent increase was observed in the EZ. Compared to N3 sleep, both the EZ and nEZ exhibited greater amplitudes in 10–200 Hz during N2 (Fig. 2B; *p* < 0.05, Wilcoxon rank sum test, multiple comparison corrected with BH procedure). The effect size of these differences (Fig. 2B, bottom) ranged from small to moderate (corresponding to Cohen’s *d* = 0.2 and 0.5, respectively).

The EZ and nEZ exhibited different bistability spectra. Compared to N3, the EZ showed smaller 5–45 Hz bistability but larger 130–225 Hz bistability during N2 (Fig. 2C). Conversely, the nEZ showed larger 2–13 Hz bistability and smaller 20–100Hz bistability during N2, with effect size ranging from small to moderate (Fig. 2C, bottom). Moreover, compared to nEZ, the EZ exhibited stronger amplitudes across slow and fast oscillations in both N2 and N3, with a large effect size (Cohen’s *d* > 0.8) in 2–30 Hz oscillations during N2 (Fig. 2D). During N2, the nEZ showed stronger 2–7 Hz bistability. EZ showed stronger bistability of high-gamma oscillations, with the largest effects in 15–150 Hz during N2 and 6–40 Hz during N3 (Fig. 2E).

In summary, the nEZ demonstrated lower amplitudes but larger bistability in δ-oscillations during N2, while the EZ showed lower δ-bistability (*i.e.*, more stable than nEZ) but increased β–γ bistability in N2.

### Strong δ and β–γ band synchrony characterizes the EZ

We examined whether large-scale phase synchrony differentiates N2 from N3 sleep and the EZ from nEZ channels. Both eigenvector centrality (EVC) and the clustering coefficient (CC) of the synchrony networks showed weak differentiation between N2 and N3 (Fig. 3A–B). However, both the EZ and nEZ exhibited stronger 2–5 Hz CC during N2 compared to N3 (Fig. 3B, bottom).

During both N2 and N3, the EZ consistently displayed stronger EVC in 2–5 Hz (Cohen’s *d* > 0.5) and 20–40 Hz (Cohen’s *d* > 0.8) synchrony, while the nEZ showed stronger EVC in 100–300 Hz synchrony, particularly during N2 (Fig. 3C). Additionally, the EZ exhibited stronger CC in the 2–5 Hz range in both N2 and N3 (Fig. 3D).

Together, these findings suggest that elevated δ and β–γ band synchrony characterize the connectivity of the EZ (characterized by EVC) and its neighboring regions (characterized by CC), with a stronger effect during N2 sleep.

### Large-scale δ–β band phase coupled with EZ’s β–γ band amplitudes (PAC) during N2

The EZ exhibits large β–γ band bistability (Fig. 2) and elevated δ and β–γ band synchrony (Fig. 3), and a potential system-level mechanistic link between these bands was next investigated using PAC analysis. We expected that β–γ amplitudes would be modulated by the δ phase, similar to what has been observed in focal epilepsy in both sleep^44^ and resting-state^40^.

Across four quadrants of the cohort-pooled PAC matrices (Fig 1F), we observed spectrally widespread connectivity in both N2 and N3 (Fig. 4A–B). In N3, the nEZ→nEZ quadrant demonstrated stronger PAC strength (*p* < 0.05, BH corrected) compared to N2 with large effect size (Cohen’s |*d*| > 0.8), particularly between the slowest δ-oscillation (< 1Hz) and β–γ amplitudes (Fig. 4C). Conversely, the EZ→EZ quadrant showed greater PAC strength during N2, most pronounced between the 15– 24Hz phase and the amplitudes of fast rhythms (Cohen’s |*d*| > 0.65). The EZ→nEZ quadrant showed greater PAC in N2 between β-phase and high-γ amplitudes with a small effect (Cohen’s |*d*|> 0.25). Most notably, in N2, the nEZ→EZ quadrant showed stronger PAC between 1–2 Hz phase and widespread fast oscillations, indicating the modulation of γ and high-γ amplitudes in the EZ by δ phase from the nEZ (Fig. 4C, right). These patterns suggest that slow δ oscillations may coordinate fast β–γ dynamics across regions, providing a potential mechanism for large-scale network interactions during interictal sleep.

### PAC modulated by δ phase is correlated with epileptogenic β–γ bistability

We next examined the hypothesis that PAC serves as a cross-frequency mechanism linking δ-phase synchrony to β–γ bistability; if so, it should correlate with both synchrony and bistability in a direction-specific manner (Fig. 5A). In this framework, outward-PAC describes how the phase of slow waves in one region modulates fast activity in other regions, whereas inward-PAC reflects how slow waves from the surrounding network modulate fast activity within that region. Rather than computing pairwise correlations independently across frequencies, as done previously, we applied CCA (Fig. 5B) to identify multivariate relationships among population-pooled PAC, synchrony, and bistability features (Fig. 5C).

We first applied CCA to synchrony and outward-PAC to confirm whether the analysis reproduces known PAC spectral characteristics, thereby validating the new approach. Synchrony and outward-PAC features were projected into canonical variates (CVs) in a latent space that maximizes their correlation, with each frequency’s contribution characterized by the loading (Fig. 5B, see Methods for details). This analysis revealed strong correlations (*r* = 0.82 for N2 and *r* = 0.80 for N3). The loadings of δ synchrony and α (7–11 Hz) and γ components of outward-PAC were the strongest contributors in both N2 and N3 (Fig. 5D). The α and γ peaks in the outward-PAC loadings matched those observed in the mean δ-modulated PAC connectivity (dashed lines, Fig. 5D), supporting the validity of the CCA results.

CCA of inward-PAC and bistability revealed strong correlations (*r* = 0.86 for N2 and *r* = 0.84 for N3), primarily driven by bistability in the 2–7 Hz and 20–100 Hz ranges, together with inward-PAC amplitudes in the 20–250 Hz range (Fig. 5E), showing stronger δ bistability involvement during N2. These results indicate that regions whose β–γ amplitudes are strongly modulated by δ-phase also exhibit β–γ bistability, supporting a cross-frequency mechanistic link between δ-phase and β–γ bistability.

### Correlation of inward-PAC and β–γ bistability with interictal spikes

Lastly, we examined the functional relevance of the canonical correlations between PAC, bistability, and synchrony by relating their canonical variates (CVs) to interictal spikes. If δ-phase and β–γ bistability are coupled through a PAC mechanism, their corresponding CVs should correlate with spike activity—an established biomarker of epileptogenicity. We averaged the CVs of channel-wise PAC, bistability, and synchrony estimates (samples shown in Fig. 5D–E scatter plots) within each subject and assessed correlations with the number of spatial spike bursts observed in the same N2 and N3 epochs. Strong correlations were found between spikes and the CVs of inward-PAC and bistability (*r*^2^=0.62 for N2 and 0.56 for N3), whereas correlations between spikes and the CVs of outward-PAC and synchrony were weak (*r*^2^=0.22 for N2 and 0.1 for N3). Note that *r^2^* values were reported because loadings were optimized to maximize correlations (Eq. 9), making correlation signs irrelevant. These findings demonstrate the functional relevance of the canonical correlations linking δ-modulated inward-PAC and β–γ bistability.

## Discussion

To investigate the systems-level mechanisms linking epilepsy-related δ- and β–γ-band oscillations, we analyzed N2 and N3 sleep in patients with SHE and underlying FCD2—a prototypical model of sleep-related epilepsy that allows accurate EZ localization and thus high-confidence comparisons between EZ and nEZ. We first replicated well-established EZ biomarkers, including δ-instability, δ-amplitude, β–γ bistability, and δ-phase to β–γ-amplitude coupling (PAC), thereby ensuring rigor and providing a foundation for the novel CCA-based analyses. Spectral–spatial unfolding of the classic δ-band instability of NREM sleep^19^ revealed novel, concurrent patterns: (1) stronger δ amplitudes in the EZ but greater δ bistability in the nEZ; (2) elevated δ- and β–γ-band connectivity involving the EZ; and (3) a strong cross-frequency coupling between δ-phase from the nEZ and β–γ amplitudes in the EZ. A strong multivariate correlation (*r* = 0.86 for N2; *r* = 0.84 for N3) between δ-modulated PAC and β–γ bistability, identified by the CCA, supports a robust cross-frequency mechanistic link between slow and fast oscillations—reflecting the ‘organic whole’ of epileptogenicity. These results support our hypothesis that bistable δ oscillations during NREM sleep modulate epileptogenic β–γ band bistability. Our results also provide strong motivation for a comprehensive EZ localization framework based on a latent space^22,34,45^ that builds on recent advances integrating multiple biomarkers from interictal NREM^46^ or resting-state^22^ SEEG.

### Analysis of local dynamics and large-scale synchrony reveals frequency-specific, EZ-specific differences between N2 and N3

In addition to the established δ-instability metric, two amplitude and two synchrony metrics were used to characterize spectral differences in local and network features between the EZ and nEZ, as well as between N2 and N3 sleep. The EZ and nEZ showed weak to moderate but distinct frequency-specific differences across sleep stages. During N3, the nEZ exhibited moderately stronger δ amplitude, reflecting the prominent slow-wave oscillations characteristic of N3. In contrast, during N2—when seizures most frequently occur in Sleep Hypermotor Epilepsy^20^—the EZ displayed stronger β-γ bistability, indicative of heightened epileptogenicity, consistent with focal epilepsy during resting-state^22^. However, synchrony differences between N2 and N3 were minimal in both the EZ and nEZ. Only during N3 did the EZ show moderate differences in the clustering coefficient for the δ phase, suggesting that its connected neighbors (both EZ and nEZ) are tightly connected. This also indicates that the elevated δ-band oscillations in the nEZ are coupled across regions.

### Strong differences between the EZ and nEZ

The EZ exhibited stronger δ-band amplitudes, particularly during N2, coinciding with pathological δ-instability in the EZ^19^. Strong β-γ band bistability identifies the EZ in both N2 and N3, suggesting a pathological role^47^ similar to that observed in focal epilepsy in resting-state^22^. Whether the downward shift of the bistability peak—from 10–300 Hz in N2 to 4–150 Hz in N3—reflects post-seizure slowing^26,48^ associated with decreased seizure risk warrants future investigation.

On a large scale, the EZ exhibited stronger phase synchrony, including δ-band EVC and CC, indicating that the EZ and its neighbors (*i.e.*, both EZ and nEZ) are strongly connected in the slow oscillations. In contrast, only stronger β-γ EVC characterized the EZ, coinciding with concurrent strong β-γ bistability within the EZ, suggesting that these pathological features in fast oscillations are confined to the EZ and not its network neighbors.

In conclusion, widespread δ synchrony and EZ-specific β–γ bistability and connectivity together point to a cross-frequency link between slow and fast oscillations.

### Novelty: strong multivariate correlations between δ-phase and β–γ bistability

Prior studies have reported only weak to moderate correlations between synchrony and criticality across narrow-band frequencies in SEEG^22,24,34^, questioning the existence of a strong mechanistic link within discrete slow and fast frequencies. Our PAC analysis also revealed moderate differences between N2 and N3. During N3, a stronger 1-Hz PAC was observed within the nEZ, indicative of a strong modulatory effect of large-scale slow oscillations on faster brain rhythms^9^. In contrast, 2–4 Hz oscillations in the nEZ strongly modulate β-γ activities in the EZ, predominantly during N2. These results imply the presence of at least two distinct δ oscillations, as previously reported in humans^49–51^ and rodent studies^52^.

In contrast, CCA revealed that δ-modulated inward-PAC was strongly correlated with bistability, indicating that regions where β-γ amplitudes were coupled to δ-phase also exhibited high β-γ bistability. We therefore speculate that enhanced δ-synchrony might therefore promote conditions that favor bistable β–γ dynamics by intermittently synchronizing local assemblies and creating alternating windows of high and low fast-activity expression—consistent with the gating role of slow oscillations observed in animal models^53,54^.

Altogether, these results suggest a link between heightened δ-phase synchrony and increased β-γ bistability mediated by cross-frequency coupling promoted during unstable N2 sleep. This supports the idea that increased instability of δ oscillations favors aberrant phase synchronization patterns that, via a cross-frequency coupling, in turn entrain a pathological network mechanism that supports the onset and spread of fast-activity bursts, particularly within the EZ.

### Slow-wave and arousal dynamics as drivers of epileptic activity

Our findings align with and expand upon previous studies^15,55^, reinforcing the concept that slow-wave instability is an inherent and fundamental feature of sleep. In epilepsy patients, this instability may be hijacked by epileptic activity, playing a crucial role in favoring the emergence of pathological dynamics^15,56^. Recent studies have further refined this perspective, showing that arousal fluctuations, regulated by noradrenergic activity, are essential for maintaining sleep continuity, sleep-related memory processes, and cerebrospinal fluid dynamics^57^. These insights, combined with advances in animal models^58,59^, offer new perspectives on the neurophysiological mechanisms underlying these dynamics, suggesting that altered arousal fluctuations may contribute to heightened cortical excitability, ultimately promoting sleep fragmentation and facilitating epileptic activity. This growing body of evidence may open new perspectives for therapeutic strategies. Beyond conventional antiseizure medications, modulating sleep physiology may represent a promising complementary approach. Pharmacological or neuromodulatory interventions targeting arousal fluctuations could help stabilize sleep, potentially mitigating epilepsy-related sleep disturbances and reducing seizure burden. Future research should explore whether optimizing sleep stability—through targeted drugs or closed-loop neuromodulation— can disrupt this pathological cycle, offering novel treatment avenues for sleep-related epilepsies.

### Scope and limitations

We studied fourteen subjects due to practical challenges and strict selection criteria, reducing the number of suitable patients. However, the small sample size was offset by strong statistical power, supported by a relatively large dataset of 190 epochs, totaling 32 hours of interictal sleep SEEG data with uninterrupted N2 and N3 recordings. Patients with well-delineated FCD-2 foci, who predominantly have seizures during sleep, served as an ideal disease model for testing our hypothesis. It remains unclear whether these features limit the generalizability of our findings, as the observed interactions may not extend to other seizure types, more complex epileptogenic networks, or broader epilepsy populations. Further investigation is needed to determine the extent to which PAC interconnects δ phase and β-γ bistability in these other contexts.

## Conclusions

During NREM sleep, δ-band phase synchrony and strong bistability in β–γ band oscillations are not isolated epileptogenic mechanisms. Instead, they are likely to work together, with large-scale slower δ waves orchestrating local β–γ bistability across large brain networks, even involving brain areas not clinically considered seizure-prone. This cross-mode interaction may be a key player in promoting seizures.

## Acknowledgments

This project is supported by: NEXTGENERATIONEU (NGEU), the Ministry of University and Research, and National Recovery and Resilience Plan project MNESYS (PE0000006; A Multiscale Integrated Approach to the Study of the Nervous System in Health and Disease; DN. 1553 11.10.2022) awarded to L.N. and G.A. A Sigrid Jusélius Foundation fellowship (210527) awarded to SHW.

## Ethics Statement

This study was approved by the ethical committee (ID: 939) of Niguarda Hospital, Milan, Italy, and was performed according to the Declaration of Helsinki. We confirm that we have read the Journal’s position on issues involved in ethical publication and affirm that this report is consistent with those guidelines.

## Patient Consent Statement

Before electrode implantation, all patients gave written informed consent for participation in research studies and for publication of the results. The patient SEEG data and clinical information were handled anonymously.

## Abbreviations

CCA: canonical correlation analysis
EZ: epileptogenic zone
FCD: focal cortical dysplasia
PAC: phase-amplitude coupling
RDP: relative δ power, an assessment of δ amplitude instability

## Author Contributions

*Conceptualization:* GA, LN, JMP, & SHW

*Funding acquisition:* GA, LN

*Methodology:* GB, GA, JMP, SHW

*Software:* GA, GB

*Formal analysis:* GB, GA

*Resources:* GA, LN

*Data curation:* GA, LN

*Visualization:* GB, GA, LN, SHW

*Original draft:* GB, LN, SHW, GA

*Writing:* GB, LN, SHW, GA

*Supervision:* GA, SHW

## Data availability

Raw data and patient information cannot be shared due to Italian governing laws and Ethical Committee restrictions. Interim results, as well as final processed data that support the findings of this study, are available from the corresponding authors upon reasonable request.

