## Supplementary Material for "Unstable Slow Oscillations Couple with Epileptogenic Fast-Rhythm Bistability in Sleep-Related Epilepsy: An SEEG Study"

### Supplementary materials for Burlando et al.

### Graphical Abstract

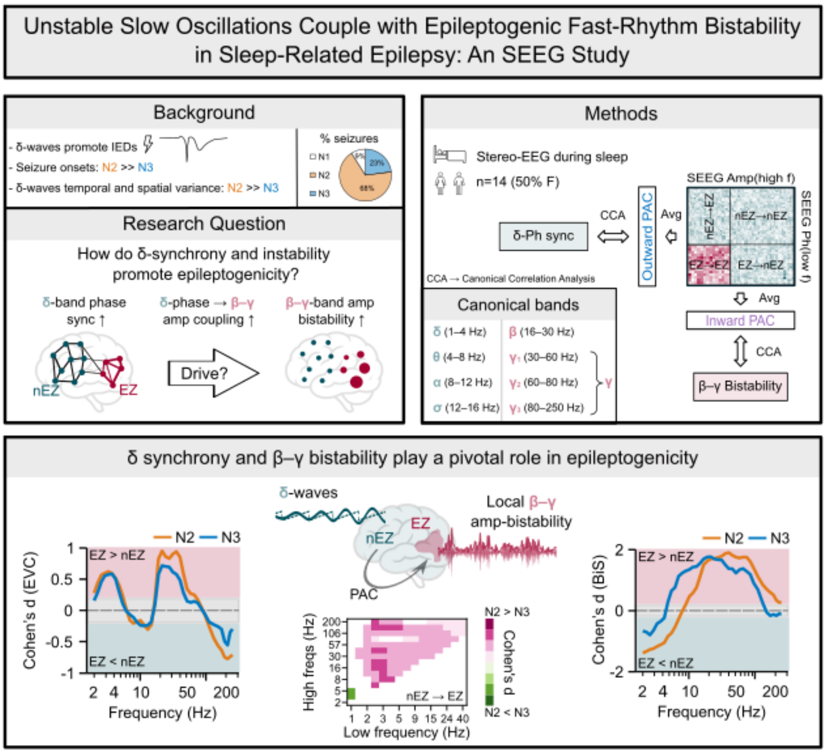


### Supplementary methods

#### Assessing features of neuronal oscillations

We followed our previous approach for assessing the δ-instability[1]. Each 10-minute epoch was divided into non-overlapping 5-second windows, yielding 120 windows. The δ-band power was defined as the average Morlet envelope between 0.5 and 4 Hz, and relative delta power (RDP) was calculated by dividing δ-power by the total power across the 0.5–180 Hz spectrum. Spatial instability was defined as the standard deviation across channels after averaging RDP over time bins, and temporal instability as the standard deviation across time bins after averaging RDP across channels.

Neuronal bistability refers to the erratic switching between an up and a down mode in a neuronal oscillation, likely caused by an underlying first-order (discontinuous) phase transition due to strong positive feedback[2,3]. The bistability index (BiS) is estimated by first fitting the sample probability distribution of the narrow-band power time series to a single-exponential (Eq. 1) and a bi-exponential model (Eq. 2), then comparing the fitting using the Bayesian information criterion (BIC). The single-exponential model is defined as:

$$P_{R^2}(R^2) = \gamma e^{-\gamma R^2} \quad (1)$$

where $\gamma$ is the exponent, and the bi-exponential model is defined as:

$$P_{R^2}(R^2) = \frac{1}{(\delta_1 + \delta_2)}(\delta_1 \gamma_1 e^{-\gamma_1 R^2} + \delta_2 \gamma_2 e^{-\gamma_2 R^2}) \quad (2)$$

where $\gamma_1$ , $\gamma_2$ are the two exponents and $\delta_1$, $\delta_2$ are weighting factors. The BIC was computed for the single- and bi-exponential fitting as follows:

$$\mathrm{BIC} = k \cdot \ln(n) - 2\ln(\hat{L}) \quad (3)$$

where $n$ is the number of samples, $\hat{L}$ is the likelihood function, $k$ is the number of free parameters in the model: $k = 1$ for single-exponential $\mathrm{BIC_{Exp}}$ and $k = 4$ for bi-exponential model $\mathrm{BIC_{biE}}$. A better-fitted model yields a small BIC value. The difference in BIC between single- and bi-exponential is obtained as:

$$\mathrm{dBIC} = \mathrm{BIC_{Exp}} - \mathrm{BIC_{biE}} \quad (4)$$

and the bistability index BiS as:

$$\mathrm{BiS} = \log_{10}(\mathrm{dBIC}) \quad \text{if } \mathrm{dBIC} > 0 \quad (5)$$

$$\mathrm{BiS} = 0 \quad \text{if } \mathrm{dBIC} \leq 0$$

A BiS close to zero indicates that the single-exponential model best fits the time series, while a BiS > 3 offers strong support for the bi-exponential model.

#### Assessing interareal phase synchrony

Synchrony between all bipolar-referenced SEEG electrode contacts was estimated using the phase-locking value (PLV)[4] for the 40 narrow-band frequencies that are evenly spaced on a log10 scale. Considering $x'(t)$ as the narrowband complex wavelet coefficients of the broadband signal $x(t)$ of a contact, the complex-valued PLV (cPLV) is:

$$\mathrm{cPLV} = \frac{1}{T}\sum_{t=1}^{T} \frac{x\prime(t)}{|x\prime(t)|} \cdot \frac{y\prime*(t)}{|y\prime(t)|} \tag{6}$$

where $T$ represents the total sample number, and * is the complex conjugate. The PLV is the absolute value of the cPLV.

To minimize bias and prevent artificially inflated synchrony estimates, we removed entries from the PLV matrices where bipolar channels shared a common contact. Each PLV matrix was treated as a graph, with contacts as nodes and pairwise synchrony as edges. To estimate first-order synchrony derivatives, we used eigenvector centrality (EVC), a measure where a node's centrality is higher if it is connected to other nodes with high centrality. Second-order synchrony derivatives were evaluated using the clustering coefficient (CC), which quantifies the proportion of a node's neighbors that are also interconnected with each other.

#### Assessing phase-amplitude coupling (PAC)

PAC refers to the modulation of fast neuronal oscillation amplitudes by the phase of slower oscillations in the brain (Fig. 1F) and was assessed using PLV[5]:

$$\mathrm{PLV}_{\mathrm{PAC},a,b} = \frac{1}{N}\left|\sum_{n=1}^{N} e^{i\cdot(\theta_{a,LF}-\theta_{b,HF,LF})}\right| \tag{7}$$

where $\theta_{a,LF}$ is the phase of signal $a$ filtered at the low frequency (LF) with a Morlet filter, while $\theta_{b,HF,LF}$ is the phase of the amplitude envelope of signal $b$, obtained by Morlet-filtering $b$ at the high frequency (HF), extracting its amplitude envelope, and filtering it again at the same LF as before, thereby capturing the LF modulation of HF amplitudes. To compute the PAC, we used a design matrix comprising ten LFs, ranging from 1.2

to 68.1 Hz, and 19 slow-to-fast frequency ratios, ranging from 1:2 to 1:100. Having thus computed PAC matrices in a low-frequency-to-ratio format, we projected them to a low-frequency-to-high-frequency space for enhanced readability and interpolated missing values. Results show normalized PAC (nPAC), where nPAC = $PLV_{PAC,observed}/PLV_{PAC,surrogate}$ (see subsection "Statistical Analyses" for surrogate creation), so that nPAC > 1 indicates PAC above the null hypothesis level.

#### Canonical correlation analysis (CCA)

Canonical correlation analysis (CCA) is a statistical procedure that identifies the maximal correlations between two datasets[6] and has been previously used for analyzing high-dimensional neuroscience data[7]. It aims to find canonical weight vectors $\mathrm{u}$ and $\mathrm{v}$ that maximize the correlations between projected data:

$$\max_{\mathrm{u,v}} \rho = \frac{\mathrm{u^T}\,\Sigma_{\mathrm{AB}}\,\mathrm{v}}{\sqrt{\mathrm{u^T}\,\Sigma_{\mathrm{AA}}\,\mathrm{u}}\sqrt{\mathrm{v^T}\,\Sigma_{\mathrm{BB}}\,\mathrm{v}}} \tag{8}$$

where $\Sigma_{\mathrm{AA}} = (\mathrm{A^T A})/\mathrm{n}$ is the covariance matrix for the data array $\mathrm{A}$ (*e.g.*, inward-PAC, size $\mathrm{n} \times \mathrm{a}$, $\mathrm{n}$ is the contact number) and $\Sigma_{\mathrm{BB}} = (\mathrm{B^T B})/\mathrm{n}$ is the covariance matrix for data array $\mathrm{B}$ (*e.g.*, bistability, size $\mathrm{n} \times \mathrm{b}$). $\Sigma_{\mathrm{AB}} = (\mathrm{A^T B})/\mathrm{n}$ is the cross-covariance of $\mathrm{A}$ and $\mathrm{B}$. Solving the generalized eigenvalue problem:

$$\begin{bmatrix} 0 & \Sigma_{\mathrm{AB}} \\ \Sigma_{\mathrm{BA}} & 0 \end{bmatrix} \begin{bmatrix} \mathrm{u} \\ \mathrm{v} \end{bmatrix} = \rho \begin{bmatrix} \Sigma_{\mathrm{AA}} & 0 \\ 0 & \Sigma_{\mathrm{BB}} \end{bmatrix} \begin{bmatrix} \mathrm{u} \\ \mathrm{v} \end{bmatrix} \tag{9}$$

yields the canonical correlation coefficients $\rho$ and the weight vectors $\mathrm{u}$ and $\mathrm{v}$. The canonical variates (CV) are computed as:

$$\mathrm{Z_A} = \mathrm{Au} \text{ and } \mathrm{Z_B} = \mathrm{Bv} \tag{10}$$

where $\mathrm{Z_A}$ and $\mathrm{Z_B}$ (both with size $\mathrm{n} \times 1$) are the CVs of $\mathrm{A}$ and $\mathrm{B}$; $\mathrm{u}\ (\mathrm{a} \times 1)$ and $\mathrm{v}\ (\mathrm{b} \times 1)$ are the first canonical weight vectors.

#### Statistical Analyses

We calculated null-hypothesis distributions for both PLV and PAC using surrogate data. For each contact pair, we split each narrow-band time series into two segments at a random time point, denoted as $\mathrm{k}$. This resulted in two time series:

$$\mathrm{x'}_1(\mathrm{t}) = \mathrm{x'}(1 \ldots \mathrm{k}) \text{ and } \mathrm{x'}_2(\mathrm{t}) = \mathrm{x'}(\mathrm{k} \ldots \mathrm{T}) \quad (11)$$

We then created the surrogate time series, $\mathrm{x'}_{\mathrm{surr}}(\mathrm{t})$, by concatenating these two segments in reverse order: $\mathrm{x'}_{\mathrm{surr}}(\mathrm{t}) = [\mathrm{x'}_2, \mathrm{x'}_1]$. The surrogate PLV and PAC were computed across all pairs of channels.

The Wilcoxon rank-sum test was used to make comparisons between groups. For PAC, the test was applied to each LF–HF pair over epochs into which the signal was segmented. For amplitude, BiS, EVC, and CC, the test was applied to each frequency across the pooled channels. Benjamini-Hochberg (BH) procedure was used to correct for multiple comparisons[8], with the post-correction significance threshold set for $p <$ 0.05. Effect size for between-group difference was evaluated with Cohen's *d*, and its confidence intervals under the null-hypothesis were measured by label-shuffled surrogates (N=10'000).

#### Validation of narrow-band differences in features

As validation of narrow-band differences in the features, we computed average effect sizes within canonical frequency bands, focusing only on those greater than an absolute value of 0.3, indicative of a medium effect size. As expected, the amplitude was well distinguished between N2 and N3 within the nEZ tissue, highlighting $\mathrm{Amp}_{N2} < \mathrm{Amp}_{N3}$ in δ, θ, and α, while $\mathrm{Amp}_{N2} > \mathrm{Amp}_{N3}$ in the higher-frequency range. In EZ, $\mathrm{Amp}_{N2} > \mathrm{Amp}_{N3}$ for frequency bands ranging from α to $\gamma_3$ (Supplementary Fig. 1A, left). The dysplasia showed higher amplitudes both for N2 (from δ to $\gamma_1$) and for N3 (from δ to α and $\gamma_1$), compared to nEZ (Supplementary Fig. 1A, right). BiS and PLV were higher in EZ than nEZ, both in N2 and N3 sleep (Supplementary Fig. 1 to B, right). Low-frequency BiS (δ and θ) could also discriminate between the two sleep stages, with $\mathrm{BiS}_{N2} > \mathrm{BiS}_{N3}$ in the healthy tissue Supplementary Fig. 1B, left). No differences between N2 and N3 sleep were observed in PLV, in both healthy and pathological tissues (Supplementary Fig. 1C, left). δ PAC nEZ → nEZ (from δ to $\gamma_3$) and EZ → nEZ (from δ to β) were higher in N3 compared to N2 and nEZ → EZ higher in N2 compared to N3 (Supplementary Fig. 1D). Squared canonical loadings of PAC and BiS were higher in N2 than in N3, for all the frequency bands but α, in which N3 > N2 (Supplementary Fig. 1E). Effect size of the difference EZ–nEZ in canonical variates of PAC and BiS was higher in N2 than N3 in absolute value (Supplementary Fig. 1F), meaning that N2 canonical variates can better discriminate between healthy and pathological tissues.

### Supplementary Tables

#### Supplementary Table 1

| Established biomarkers | | | Novel biomarkers: Interictal resting-state SEEG | | | | | | | | |
|---|---|---|---|---|---|---|---|---|---|---|---|
| | N2 | N3 | | δ | θ | α | σ | β | $\gamma_1$ | $\gamma_2$ | $\gamma_3$ |
| δ-instability | ☑ | | LRTCs | ☑ | ☑ | | | | | | |
| δ-oscillations | | ☑ | Bistability ↑ | | | | | ☑ | ☑ | ☑ | |
| K-complexes | ☑ | | Inhibition | | | | | ☑ | ☑ | ☑ | |
| IEDs | ☑ | | 1:1 PLV | ☑ | ☑ | ☑ | | | | | |
| Seizures | ☑ | | n:m PAC | EZ–EZ: Θ(δ–α) → A(σ–$\gamma_3$) ↑ | | | | | | | |

***Established and novel biomarkers proposed in prior art.*** IEDs: interictal epileptiform discharges; PLV: phase-locking value; LRTCs: power-law scaling in long-range temporal correlations; PAC: cross-frequency phase-amplitude coupling. EZ: epileptogenic zone. Canonical bands: δ (1–4 Hz), θ (4–8 Hz), α (8–12 Hz), σ (12–16 Hz), β (16–30 Hz), $\gamma_1$ (30–60 Hz), $\gamma_2$ (60–80 Hz), $\gamma_3$ (80–250 Hz), γ → $\gamma_{1\text{-}3}$.

#### Supplementary Table 2

| Subject | Explored Anatomical Areas | Epileptogenic Zone | FCD type |
|---|---|---|---|
| **01** | Frontal, Central, Parietal, Operculo-insular (R) | Operculo-insular (R) | IIa |
| **02** | Frontal, Central, Parietal, Operculo-insular (R) | Cingulate (R) | IIa |
| **03** | Frontal, Central, Parietal, Temporal, Operculo-insular (R) | Insular (R) | IIa |
| **04** | Frontal, Temporal, Operculo-insular (R) | Orbital (R) | IIb |
| **05** | Frontal, Central, Parietal, Operculo-insular (L) | Operculo-insular (L) | IIb |
| **06** | Frontal, Operculo-insular (R) | Mesial frontal (R) | IIa |
| **07** | Frontal, Temporal (R) | Fronto-orbital (R) | IIa |
| **08** | Frontal, Central, Parietal (bilateral) | Fronto-orbital (R) | IIb |
| **09** | Frontal, Operculo-insular (bilateral) | Frontal (R) | IIb |
| **10** | Frontal, Central, Operculo-insular (bilateral) | Frontal (R) | IIa |
| **11** | Frontal, Central, Parietal (bilateral) | Cingulate frontal (L) | IIb |
| **12** | Frontal, Central, Parietal, Temporal (R) | Parietal (Precuneus, R) | IIb |
| **13** | Frontal (L) | Frontal (L) | IIb |
| **14** | Frontal, Central, Operculo-insular (L) | Frontal (L) | IIb |

***Patient-wise summary of SEEG exploration.*** For each patient, the brain regions explored during SEEG implantation, the epileptogenic zone (EZ), and the type of cortical dysplasia are reported.

### Supplementary Figures

#### Supplementary Figure 1

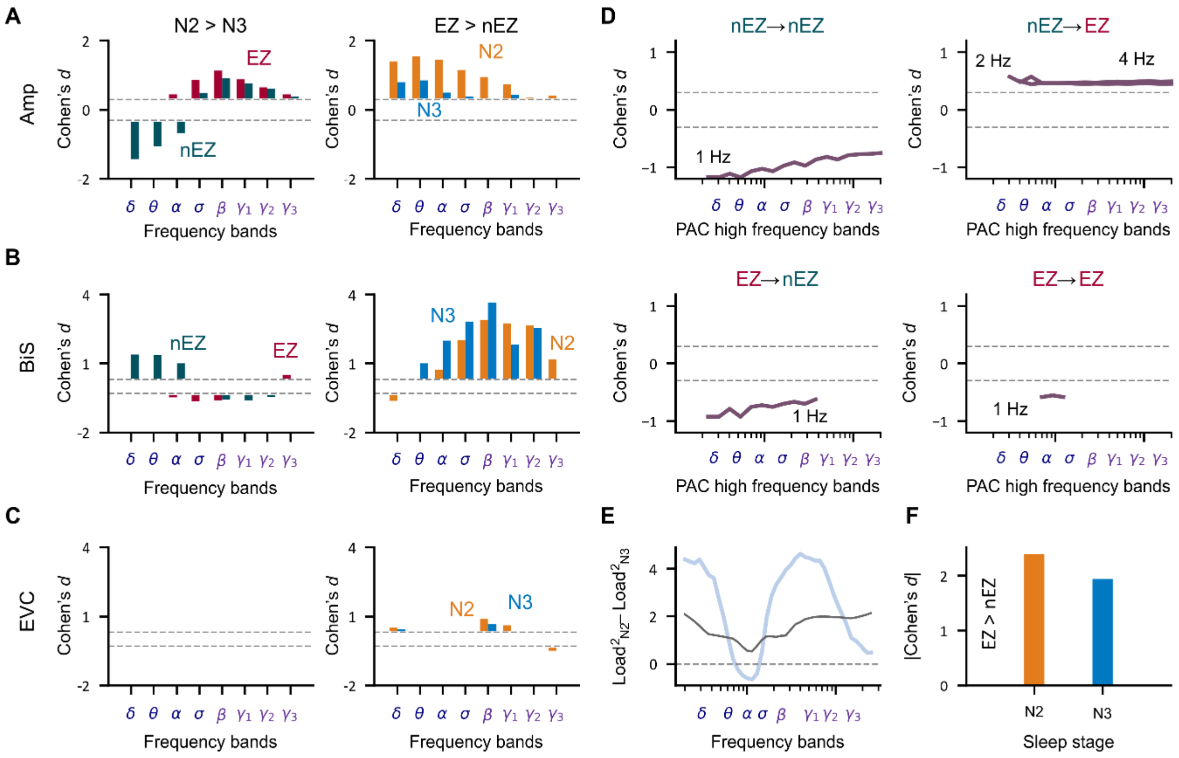


***Overview of differences.*** **(A)** Averaged Cohen's *d* within each canonical frequency band for differences in amplitude, **(B)** in BiS, and **(C)** in EVC. Only |*d*| > 0.3 are shown, indicating a small/medium effect. On the left, positive values indicate N2 > N3, while on the right indicate EZ > nEZ. **(D)** Cohen's *d* values for differences in delta PAC between N2 and N3. Positive values indicate N2 > N3. **(E)** Differences in squared canonical loadings for delta PAC and BiS. The PAC difference was obtained by interpolating the three different delta PAC lines. **(F)** Absolute effect size of canonical variates for PAC-BiS in N2 and N3.

### Supplementary Figure 2

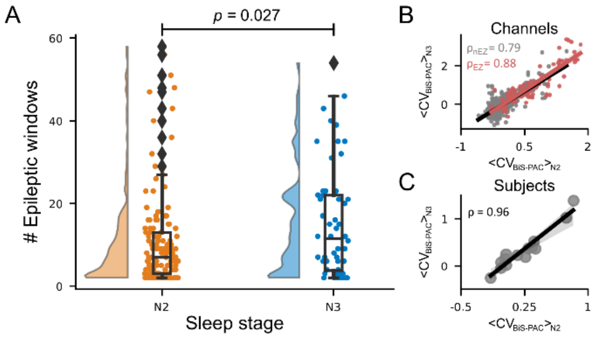


***Spike count and correlations.*** **(A)** Raincloud plot showing the distribution of the number of spiky windows during N2 and N3. Each dot represents a session ($N_{N2}$ = 134, $N_{N3}$ = 56). **(B–C)** Pearson's correlations (ρ) between canonical variates (CVs) of inward-PAC and bistability, analyzed separately for **(B)** different channels and **(C)** each subject.

### Supplementary Figure 3

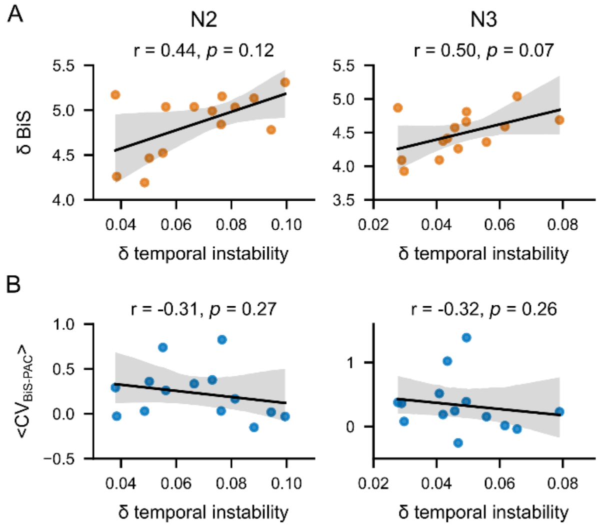


***Positive correlation arises between delta bistability and temporal instability.*** **(A)** Scatter plot showing the correlation between delta bistability (delta BiS) and delta temporal instability. **(B)** Scatter plot showing the correlation between canonical variates (CVs) of inward-PAC and bistability and delta temporal instability. For both (A) and (B), Spearman's correlation coefficient (r) and the corresponding *p*-value are reported.

#### Supplementary Figure 4

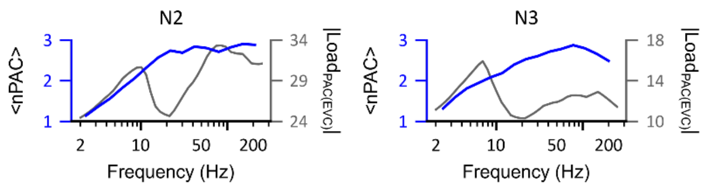


***Delta normalized phase-amplitude coupling (nPAC) and PAC loadings derived from CCA with EVC exhibit similar spectral profiles.*** Spectra of delta all-to-all nPAC (blue lines) and CCA-derived PAC loadings (grey lines) are shown for stage N2 (left) and stage N3 (right).
